# The initiation and early development of the tubulin-containing cytoskeleton in the human parasite *Toxoplasma gondii*

**DOI:** 10.1101/2023.11.03.565597

**Authors:** Luisa F. Arias Padilla, John M. Murray, Ke Hu

**Author notes:** Address correspondence to:* Ke Hu.

## Abstract

The tubulin-containing cytoskeleton of the human parasite *Toxoplasma gondii* includes several distinct structures: the conoid, formed of 14 ribbon-like tubulin polymers, and the array of 22 cortical microtubules (MTs) rooted in the apical polar ring. Here we analyze the structure of developing daughter parasites using both 3D-SIM and expansion microscopy. Cortical MTs and the conoid start to develop almost simultaneously, but from distinct precursors near the centrioles. Cortical MTs are initiated in a fixed sequence, starting around the periphery of a short arc that extends to become a complete circle. The conoid also develops from an open arc into a full circle, with a fixed spatial relationship to the centrioles. The patterning of the MT array starts from a “blueprint” with ∼ 5-fold symmetry, switching to 22-fold rotational symmetry in the final product, revealing a major structural rearrangement during daughter growth. The number of MT is essentially invariant in the wild-type array, but is perturbed by the loss of some structural components of the apical polar ring. This study provides insights into the development of tubulin-containing structures that diverge from conventional models, insights that are critical for understanding the evolutionary paths leading to construction and divergence of cytoskeletal frameworks.

## INTRODUCTION

The cytoskeleton provides the structural basis underlying cell shape and mechanical stability, as well as various intracellular activities such as vesicular transport and force generation. The characteristic cytoskeletal arrangement in different cell types is coupled with specific cellular function and activities. Among the thousands of species of apicomplexan parasites, one of the shared characteristic cytoskeletal architectures is the set of cortex-associating (variously denoted as “subpellicular” or “cortical”) microtubules (MTs) (Figure 1). The number of MTs and their arrangement within the array is species-specific, and highly reproducible within each species (Garnham *et al*., 1962; Bannister and Mitchell, 1995; Morrissette and Sibley, 2002a; Bounkeua *et al*., 2010; Wang *et al*., 2020). Apicomplexans are responsible for many high-impact and often devastating diseases, including malaria, cryptosporidiosis, and toxoplasmosis (Torgerson and Mastroiacovo, 2013; Checkley *et al*., 2015; Seeber and Steinfelder, 2016; World-Health-Organization, 2022). Their tubulin-containing cytoskeleton provides a potential rich source of drug targets (Morrissette, 2015). In *Toxoplasma gondii*, a prototypical apicomplexan, the cortical MTs are stabilized by a suite of novel associated proteins (Tran *et al*., 2012; Liu *et al*., 2013; Liu *et al*., 2015; Leung *et al*., 2017). Chemical inhibition of the elongation of the cortical MTs during cell division results in a distorted cortex and non-viable progeny (Stokkermans *et al*., 1996; Morrissette and Sibley, 2002b; Fennell *et al*., 2006; Hu *et al*., 2006; Hu, 2008). In addition, unlike most polarized microtubule-arrays examined so far, which typically originate from a centrosome or spindle pole, the cortical MTs of *Toxoplasma* and other apicomplexans are anchored in the apical polar ring, an annular structure at the parasite apex (Nichols and Chiappino, 1987; Bannister and Mitchell, 1995; Morrissette *et al*., 1997; Morrissette and Sibley, 2002a; Leung *et al*., 2017). Genetic manipulations that disrupt the integrity of the apical polar ring result in abnormalities in the MT array, failed mosquito transmission for *Plasmodium* ookinete (Qian *et al*., 2022) and highly impaired lytic cycle for the *Toxoplasma* tachyzoite (Leung *et al*., 2017; Tosetti *et al*., 2020; Back *et al*., 2021). In *Toxoplasma*, the same tubulin subunits that polymerize into the cortical MTs are also the building blocks for the conoid, which is formed of 14 ribbon-like, novel tubulin polymers organized into a left-handed spiral (Hu *et al*., 2002b; Sun *et al*., 2022; Gui *et al*., 2023; Li *et al*., 2023) (Figure 1). Like a piston, the conoid can protrude and retract in a calcium-dependent manner, passing through the apical polar ring (Hu *et al*., 2002b; Hu *et al*., 2006). The disruption of the conoid structure severely impairs parasite invasion into the host cell (Nagayasu *et al*., 2016; Long *et al*., 2017; Back *et al*., 2020; Tosetti *et al*., 2020; Dos Santos Pacheco *et al*., 2021; O’Shaughnessy *et al*., 2023).

**Figure 1.**
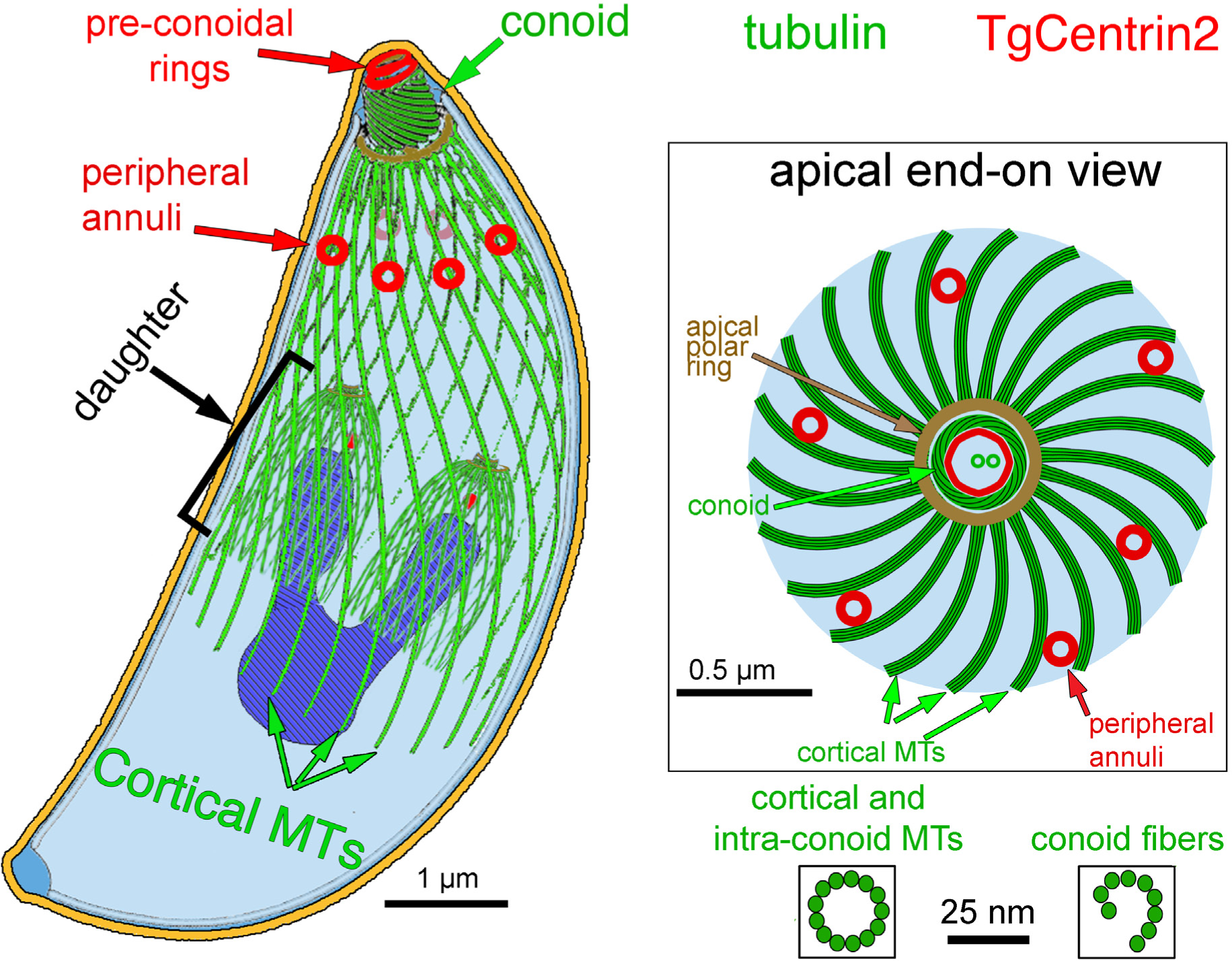
Cartoon drawings of *Toxoplasma gondii* viewed from the side (*left*) and end-on from the apical end (*right*). Tubulin-containing structures are green, TgCentrin2-containing structures are red. In the side view, the pre-conoidal rings are shown atop the conoid, and centrin2-containing annuli are distributed around the periphery. The conoid fibers are ribbon-like tubulin polymers with a protofilament arrangement very different from that in a microtubule (cross-sections shown on the lower right side). The plasma membrane is shown in orange, and the apical polar ring in brown. Inside the mature parasite, the developing cortical microtubule arrays that envelop two mid-stage daughters are shown. The nucleus (dark blue) is shown being partitioned between the daughters. Within the daughters, a red dot indicates the location of the centrin2-containing centrioles. For clarity, other organelles are not shown. Viewed from the apical end, the 22 cortical MTs are shown anchored in the apical polar ring, which encircles the conoid. The two intra-conoid microtubules (green) are seen through the pre-conoidal rings (red).

The apicomplexans use a “born-within” mode of cell division (endodyogeny, endopolygeny, or schizogony). During cell division, the daughter cytoskeleton is built from scratch instead of rearranged from the pre-existing maternal cytoskeleton, which allows immediate and unambiguous recognition of the daughter cytoskeletons under-construction vs the fully-established maternal structure (Hepler *et al*., 1966; Sheffield and Melton, 1968; Sheffield, 1970; Dubremetz and Elsner, 1979; Hu *et al*., 2002a; Morrissette and Sibley, 2002a; Voß *et al*., 2023). The co-existence of mature (*i.e.*, mother) as well as forming (*i.e.*, daughter) cytoskeletons also provides a convenient model to address how distinct biochemical states of apparently identical cellular structures can exist in the same cell within rapid diffusion distance from each other, and how one state can develop into the other (Liu *et al*., 2015; Tengganu *et al*., 2023). Together, the stunning reproducibility, ordered arrangement, and readily observed *de novo* construction of the tubulin-containing cytoskeleton in apicomplexans enable the discovery of structural features and transitions difficult to detect in other less well-ordered systems. It is therefore a superior model for determining the structural basis for constructing a defined cytoskeletal framework.

The *Toxoplasma gondii* tachyzoite has 22 spirally arranged cortical MTs (Figure 1). In mature parasites, these MTs form an ultra-stable “rib cage” and remain intact indefinitely (in experimental terms) under treatments that depolymerize MTs in mammalian cells within minutes (Morrissette *et al*., 1997; Hu *et al*., 2002b). When three associated proteins are all removed, the cortical MTs in mature parasites are destabilized, which results in surprisingly little change in parasite shape, but in a dramatic change from helical to linear parasite movement in a 3-D matrix (Liu *et al*., 2015; Tengganu *et al*., 2023). The roots of the cortical MTs are located at evenly spaced intervals around the circumference of the apical polar ring, separated by cogwheel-like projections (Nichols and Chiappino, 1987; Morrissette *et al*., 1997; Morrissette and Sibley, 2002a; Leung *et al*., 2017). Although it does not contain γ-tubulin (Morrissette, 2015; Suvorova *et al*., 2015), the apical polar ring is likely to be the organizing center for the cortical MTs. We previously showed that the cortical MTs grow at their distal ends (Hu *et al*., 2002b), suggesting that their minus ends are anchored at the apical polar ring as proposed earlier (Russell and Burns, 1984). This polarity arrangement was later confirmed by cryo-electron microscopy studies (Wang *et al*., 2021; Sun *et al*., 2022; Li *et al*., 2023). During cell division, while the mitotic spindle is assembled in the nucleus, the cortical microtubules of the developing daughter polymerize in the cytosol (Liu *et al*., 2015). Simultaneously the conoid also develops in close proximity to the apical ends of the cortical MTs (Hu *et al*., 2006; Hu, 2008).

While much progress has been made in the structural analysis of the cytoskeleton of mature parasites (Morrissette *et al*., 1997; Hu *et al*., 2002b; Hu *et al*., 2006; Wang *et al*., 2021; Sun *et al*., 2022; Gui *et al*., 2023; Li *et al*., 2023), knowledge of the initial stages of daughter development is lacking. For example, is there a defined pattern in the initial assembly process? Are all 22 cortical MTs nucleated at the same time or sequentially? Are the cytoplasmic tubulin-containing structures-the cortical MTs, conoid, and intra-conoid MTs-nucleated from the same or distinct precursor structures? Answering these questions will provide important insights into how complex supramolecular assemblies are built with structure that is identical in every cell. Here we use three-dimensional structured illumination microscopy (3D-SIM) and Expansion Microscopy (ExM) to determine the fine structural details of nascent daughters, centered around the construction of the cortical MT array and the conoid.

## RESULTS

In the mature parasite, the cortical MTs are evenly distributed around the cortex with approximate 22-fold rotational symmetry (Figure 2A). To establish the sequence of structural transitions during daughter construction in live parasites, we carried out 3D-SIM imaging of parasites expressing eGFP tagged *Toxoplasma* α1-tubulin (TUBA1) (Figure 2B-F). Nascent daughters first appear as two disc-like structures (Figure 2C), and then develop into a structure resembling an imperfect 5-petaled flower [Figure 2D, also reported in (Liu *et al*., 2015; Nagayasu *et al*., 2016; Li *et al*., 2023)], before elongating and broadening to assume a dome-like shape more similar to that of the mother parasite (Figure 2E-F). This immediately raises several questions highly relevant to the mechanism of patterning. At the front and center is: how are the cortical MTs arranged in the nascent, disc-like array precursor? Also remaining unknown is how the initiation and development of the tubulin-containing fibers of the conoid are related to that of the MT array.

**Figure 2.**
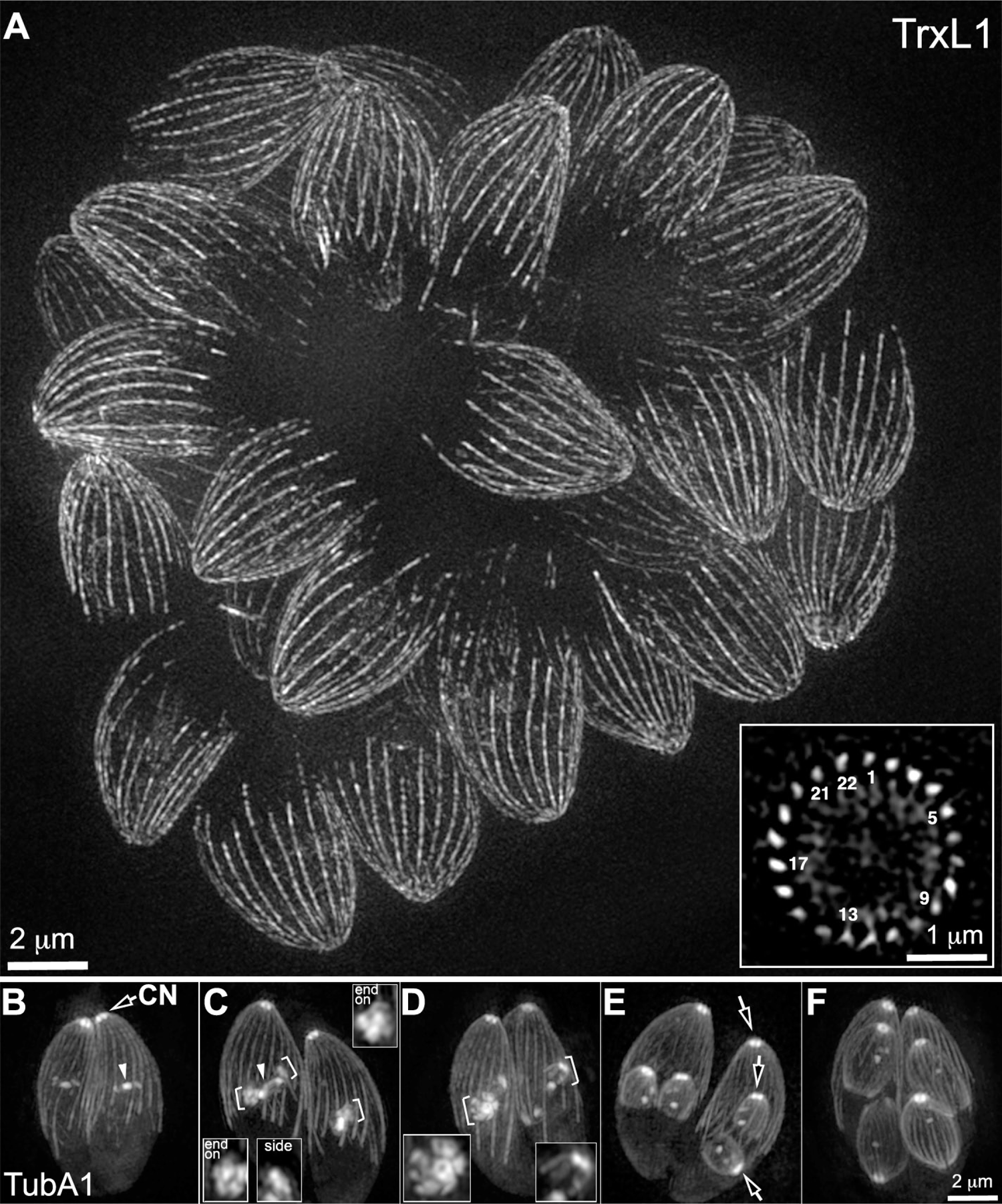
The cortical MT array switches from approximate 5-fold to 22-fold rotational symmetry during development. **A.** Projections of 3D-SIM images of live *Toxoplasma* parasites expressing mEmeraldFP-TrxL1 (Liu *et al*., 2013), in which cortical MTs are fluorescently labeled. Inset: an end-on view of an optical section through the middle of an adult parasite showing the even distribution of the 22 cortical MTs around the cortex. A subset of the MTs are labeled with numbers. **B-F.** Projections of 3D-SIM images of several live *Toxoplasma* parasites expressing eGFP-α1-tubulin at different stages of cell division. Insets (2X, contrast enhanced) include end-on or side views of the nascent daughters indicated by brackets. Image contrast was adjusted to enable better visualization of structures with weaker fluorescence (*e.g.*, cortical MTs) without oversaturating the brighter regions (e.g. conoid labeling). Arrowheads: spindles. Arrows: conoid (CN).

While 3D-SIM imaging of live parasites uncovered some unexpected structural intermediates of the growing cytoskeleton, the resolution in SIM images is insufficient to resolve fine structural details in very early nascent daughters. By expanding biological samples in a hydrogel, <u>Ex</u>pansion <u>M</u>icroscopy (ExM) has enabled the visualization of structural details much smaller than the resolution limit of conventional light microscopy (Chen *et al*., 2015; Gambarotto *et al*., 2019; Wassie *et al*., 2019; Damstra *et al*., 2022; Klimas *et al*., 2023). The ExM method (Gambarotto *et al*., 2019) has been applied in investigating subcellular details of apicomplexans [reviewed in (Liffner and Absalon, 2023)]. Here we use a slightly modified version of this method (see Materials and Methods) to determine the arrangement of tubulin containing structures in developing daughters. Our goal was to determine 1) the symmetry of the nascent MT array, *i.e.*, whether it has 5, 22-fold or other symmetry and 2) the number of MTs in the array, *i.e.*, whether all 22 MTs are present from the beginning or added incrementally. By comparing the measured diameter of the conoid in SIM images of live parasites with its diameter in ExM images, we determined that the expansion ratio in our experiments varies from ∼4.6 to 5.7 fold, with a weighted average of 5.4 fold. This was sufficient to reveal a wealth of structural detail not seen before in developing daughters. In what follows, all distance measurements made on ExM images are reported after correcting for the experimentally measured gel expansion, *i.e*., as they would have been measured in images of a live parasite, had that been possible.

Prior to the appearance of the daughter tubulin cytoskeleton, *Toxoplasma* centrioles duplicate into two pairs located adjacent to the two poles of the nascent spindle, consistent with previous findings (Hu *et al*., 2006; Hu, 2008; Francia *et al*., 2012; Chen and Gubbels, 2013; Francia and Striepen, 2014; Liu *et al*., 2015; Suvorova *et al*., 2015). The two centrioles within each pair are clearly resolved in the ExM images. When viewed end on, each centriole can be discerned as a circle (Figure 3A). While the nine singlet microtubules (Morrissette and Sibley, 2002b; Li *et al*., 2023) are not quite resolved, a substructure with 4 to 5 distinct nodes is often seen. The daughter MT arrays first initiate close to the duplicated centrioles (Figure 3B-C), as 6 to 7 “stubs” of MTs encircling the nascent conoid in a formation that resembles an incomplete smaller version of the “cogwheel” appearance characteristic of the adult apical polar ring as seen in previous EM studies (Leung *et al*., 2020). This form of structural intermediate was previously unknown and clearly demonstrates that the 22 cortical MTs are not all initiated at the same time. This also indicates that the initial incorporation of nucleation factors or activation of nucleation activity is differentially regulated across/around the MT organizing center. In the disc-like nascent daughters (Figure 3D), individual MTs can be resolved in groups of 4 and 6. The full complement of twenty-two MTs is unequivocally present when the MTs are only ∼ 0.23 µm long, *i.e.,* when there are only ∼ 30 α-β-tubulin heterodimers in each MT protofilament (Figure 3E). Thus nucleation of the cortical MTs is completed very early during daughter development. At a slightly later stage (Figure 3F), the approximate 5-fold symmetry becomes more apparent, with the MTs grouped into “petals” with 4, 4, 4, 4, and 6 MTs, consistent with the 5-petal arrangement observed in live parasites by 3D-SIM. To determine the fidelity of the number and organization of MTs in the array, we closely examined 50 very early daughter parasites in which all the cortical MTs could be resolved unambiguously. All fifty of them had 22 MTs. Twenty eight had the grouping pattern of 4, 4, 4, 4, and 6 MTs, and 15 had a pattern that could be easily derived from that grouping (*e.g.*, 4, 4, 6, 2, 6). Of note, a 4, 4, 4, 4, 4, 2 grouping was reported in a previous cryo electron-tomography study (Li *et al*., 2023). We thus propose that the grouping into 4, 4, 4, 4, and 6 MTs is the foundational pattern defined by the location of the nucleation sites.

**Figure 3.**
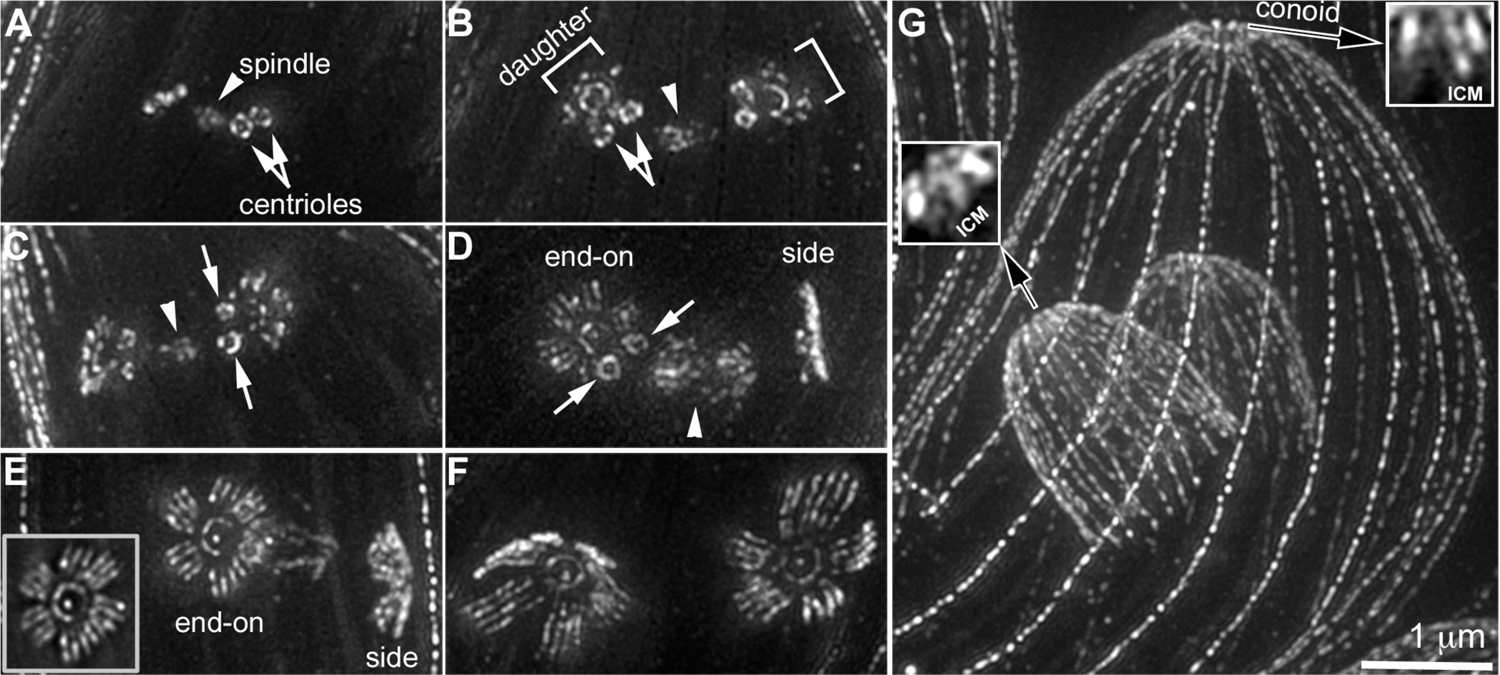
Projections of ExM images of wild-type (WT) parasites labeled with an anti-tubulin antibody. Images showing duplicated centrioles adjacent to a forming spindle prior to the initiation of daughters (A) and pairs of nascent daughters progressing from the initiation to the elongation stage (B-G). The two daughters in the pair in D and E are roughly perpendicular to each other, providing end-on and side views. The inset (1X) in E is the projection of a subset of the sections showing that all 22 MTs have been nucleated in this daughter. The approximate five-fold rotational symmetry is evident in daughters shown in F. The cortical MTs are symmetrically distributed in daughters included in G. Insets (2X, contrast enhanced) in G include a section through the conoid region of the mother (right) and a daughter parasite (left). ICM: Intra-conoid microtubules. Arrowheads: spindle. White arrows: centrioles. Brackets: initiating daughters. Black arrows: conoid. Bars ≈ 1 µm prior to expansion.

The number and length of cortical MTs in the array allow one to arrange images of early developing daughters into the correct chronological order (Figure 4). Close examination of the daughter anlagen shows that the conoid co-develops in close proximity to the cortical MT array. The end-on view of the first detectable conoid precursor shows a ∼ 0.22 μm arc (Figure 4A), which is slightly shorter than the expected apparent length of a single conoid fiber tilted as it would be in an adult conoid (∼ 0.36 μm in length, viewed at ∼ 30° = 0.31 μm) (Leung *et al*., 2020). The arc therefore represents only a few, perhaps only one, polymerizing conoid fibers.

**Figure 4.**
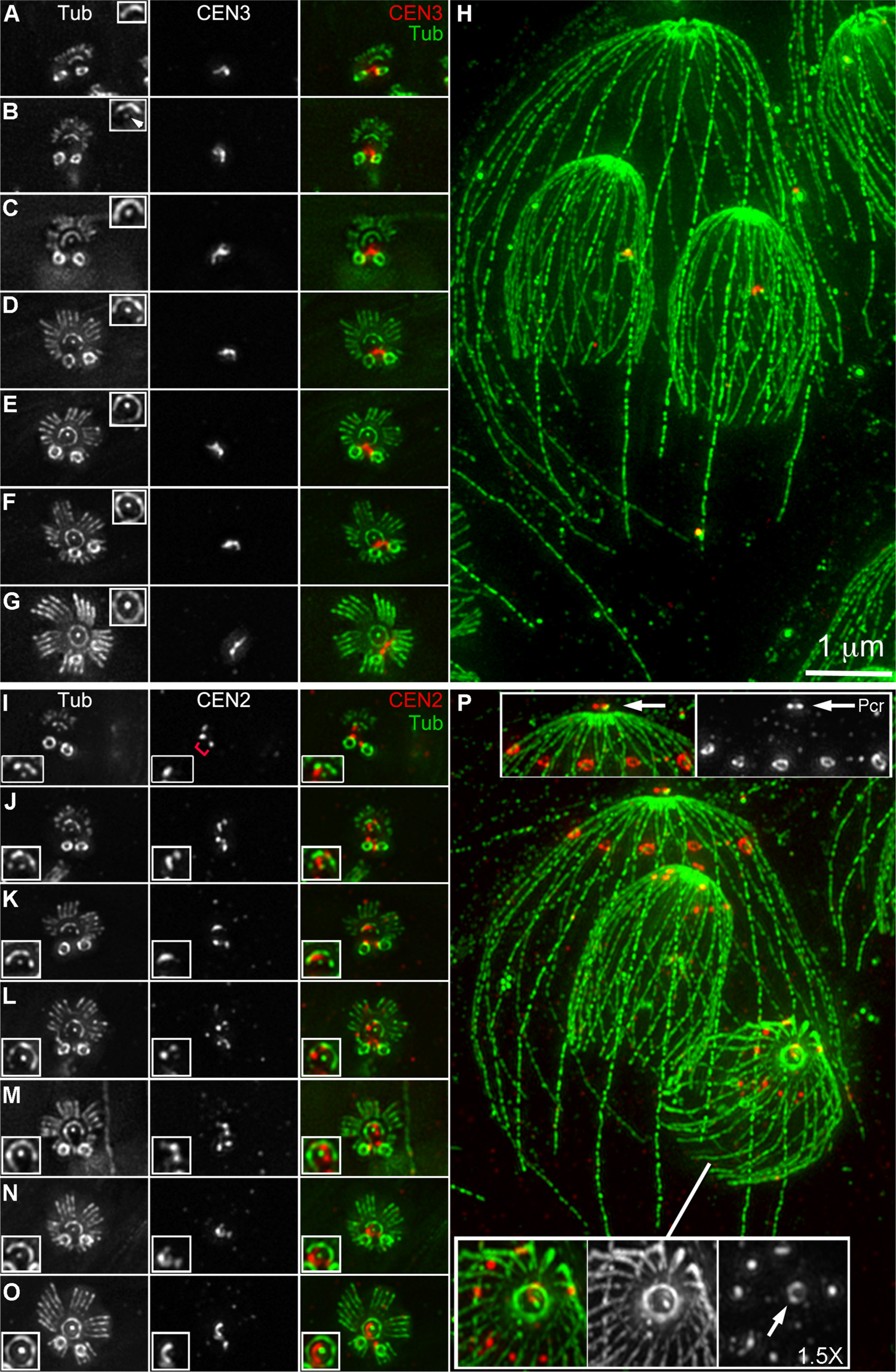
The cortical MT array and the conoid in the daughter anlagen grow towards the centrioles. **A-H**: Projections of ExM images of WT parasites labeled with an anti-tubulin (gray and green) and an anti-CEN3 antibody (gray and red). A-G: nascent daughters progressing from initiation to the stage at which the approximate five fold rotational symmetry has been established. Insets (1.5X, contrast enhanced) include the developing conoid with the intra-conoid MTs close to the center (white arrowhead in the inset in B). H: A dividing parasite in which cortical MTs are symmetrically distributed in daughters. **I-P**: Projections of ExM images of WT parasites labeled with an anti-tubulin (gray and green) and an anti-CEN2 antibody (gray and red) with the same arrangement as A-H. In addition to centrioles, CEN2 highlights the <u>p</u>re-<u>c</u>onoidal <u>r</u>ings (Pcr) (indicated by arrows in insets in P). The CEN2 signal in the pre-conoidal rings initially appears as a small focus or a short arc (insets, I-O) adjacent to the intra-conoid MTs. CEN2 also highlights the peripheral annuli [top insets (1X) in P]. Note that the peripheral annuli are not detected in early daughters (I-O), but are clearly present in later stage daughters where MTs are more evenly distributed (P, bottom insets, 1.5X, contrast enhanced). Bars ≈ 1 µm prior to expansion.

The intra-conoid MTs are polymerized slightly after the conoid and cortical MT precursors can be first detected (Figure 4B). In the end-on view, the assembly of the conoid and the cortical MT array always proceed in the same direction, forming arcs with their open side (i.e., the yet-to-be assembled side) oriented towards the centrioles (Figure 4A-E). The two centrioles are connected by centrin fibers throughout daughter development, with the fibers invariably interacting with the conoid-facing side of the centrioles, as indicated by the labeling with an anti-centrin3 (CEN3) antibody (Figure 4A-H).

In mature parasites, the conoid fibers are attached to a set of ring-like structures (the pre-conoidal rings, Fig 1) at their apical ends (Morrissette *et al*., 1997; Hu *et al*., 2002b; Sun *et al*., 2022; Gui *et al*., 2023). We previously found that another *Toxoplasma* centrin, TgCentrin2 (CEN2), is localized to not only the centriole but also the pre-conoidal rings, as well as in 5-6 peripheral annuli (Hu *et al*., 2006; Leung *et al*., 2019). Here we examined how these centrin-based structures develop with respect to each other and to the tubulin-containing structures (Figure 4I-P). We found that similar to CEN3, CEN2 persists at the centrioles throughout daughter development. However, rather than connecting the two centrioles, CEN2 often appears as two distinct foci (Figure 4I, red bracket). In late daughters or adult parasites, the pre-conoidal CEN2 is organized into a complete ring (Figure 4P). In nascent daughters, the pre-conoidal CEN2 signal first appears as a small focus in the immediate neighborhood of the nascent conoid (Figure 4I). Even when the conoid appears to be a complete circle in the end-on view, the pre-conoidal CEN2 signal is not a ring but an arc, indicating that the pre-conoidal rings are not yet established at this point (Figure 4O). In view of their tight association with the conoid fibers (Morrissette *et al*., 1997; Hu *et al*., 2002b; Hu *et al*., 2006; Munera Lopez *et al*., 2022; Sun *et al*., 2022; Gui *et al*., 2023), the pre-conoidal rings might be the organizing center for the conoid. However, this lag between conoid and pre-conoidal ring maturation seems to argue for some other nucleation sites for conoid fibers.

In adult parasites, the peripheral CEN2 signal is resolved in ExM images as annuli (Figure 4P), consistent with the previous immunoEM analysis (Hu *et al*., 2006). In daughter parasites, the peripheral CEN2 structures are first detected as spots slightly larger than the point spread function separated by two, four or six MTs (Figure 4P). These spots appear long after the 5-petaled MT array is fully established, indicating that although the CEN2 annuli also display approximate 5-fold symmetry, they do not play a role in influencing the structure of the nascent daughter MT array.

The cortical MTs are connected to the apical polar ring, a putative organizing center of the MT array. Previously we discovered that kinesinA and APR1 are two components of the apical polar ring (Leung *et al*., 2017). The double knockout of the two genes results in the displacement of the cortical MTs from the apex of adult parasites due to the disintegration of the apical polar ring. To determine how and when the structural perturbation of the MT array occurs, we examined the MT structure of daughter parasites of the *ΔkinesinAΔapr1* mutant. Upon establishment of the MT petals, 100% (50/50) of the WT daughters have 22 cortical MTs (Figure 5A). However, ∼44 % (12/27) of daughter parasites in the *ΔkinesinAΔapr1* mutant have 23 or 24 MTs (Figure 5B). This indicates that while the nucleating activity *per se* for cortical MTs is not affected by the removal of kinesin A and APR1, the number of nucleation sites becomes more variable without these scaffold proteins, which affects the fidelity of assembly of the MT array. Aside from the variable number of MTs, the structure of the early daughter MT array in the *ΔkinesinAΔapr1* mutant appears normal, even though MT detachment is pronounced in the mother parasite (Figure 5B and Figure 6A). This indicates that factors other than or in addition to kinesinA and APR1 are reponsible for MT attachment in early daughter development.

**Figure 5.**
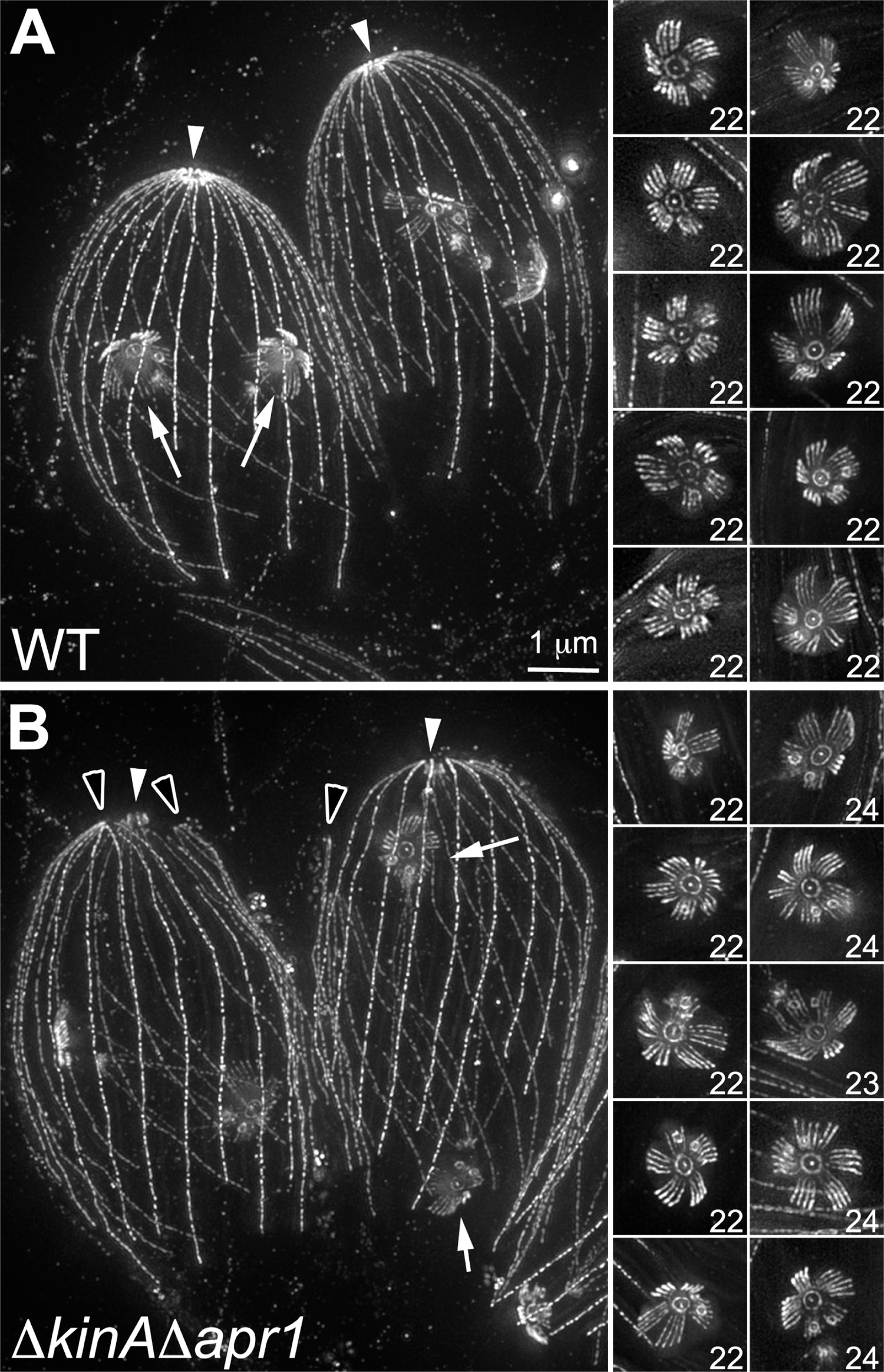
The loss of Kinesin A and APR1, two apical polar ring components, results in variation in MT numbers in the nascent array. Projections of ExM images of WT (A) and *ΔkinesinAΔapr1* parasites (B) labeled with an anti-tubulin antibody. A and B include two dividing WT (A) or *ΔkinesinAΔapr1* (B) parasites with early daughters. Note that clusters of MTs are detached from the apex of the mature *ΔkinesinAΔapr1* parasites (B, black arrowheads). The right panels include WT (A) or *ΔkinesinAΔapr1* (B) daughter parasites in which the MT petals are fully established. While the WT parasites always have 22 MTs, the number of MTs in the array in the *ΔkinesinAΔapr1* parasite varies between 22 and 24. Bars ≈ 1 µm prior to expansion. Arrows: daughters. White arrowheads: conoid.

**Figure 6.**
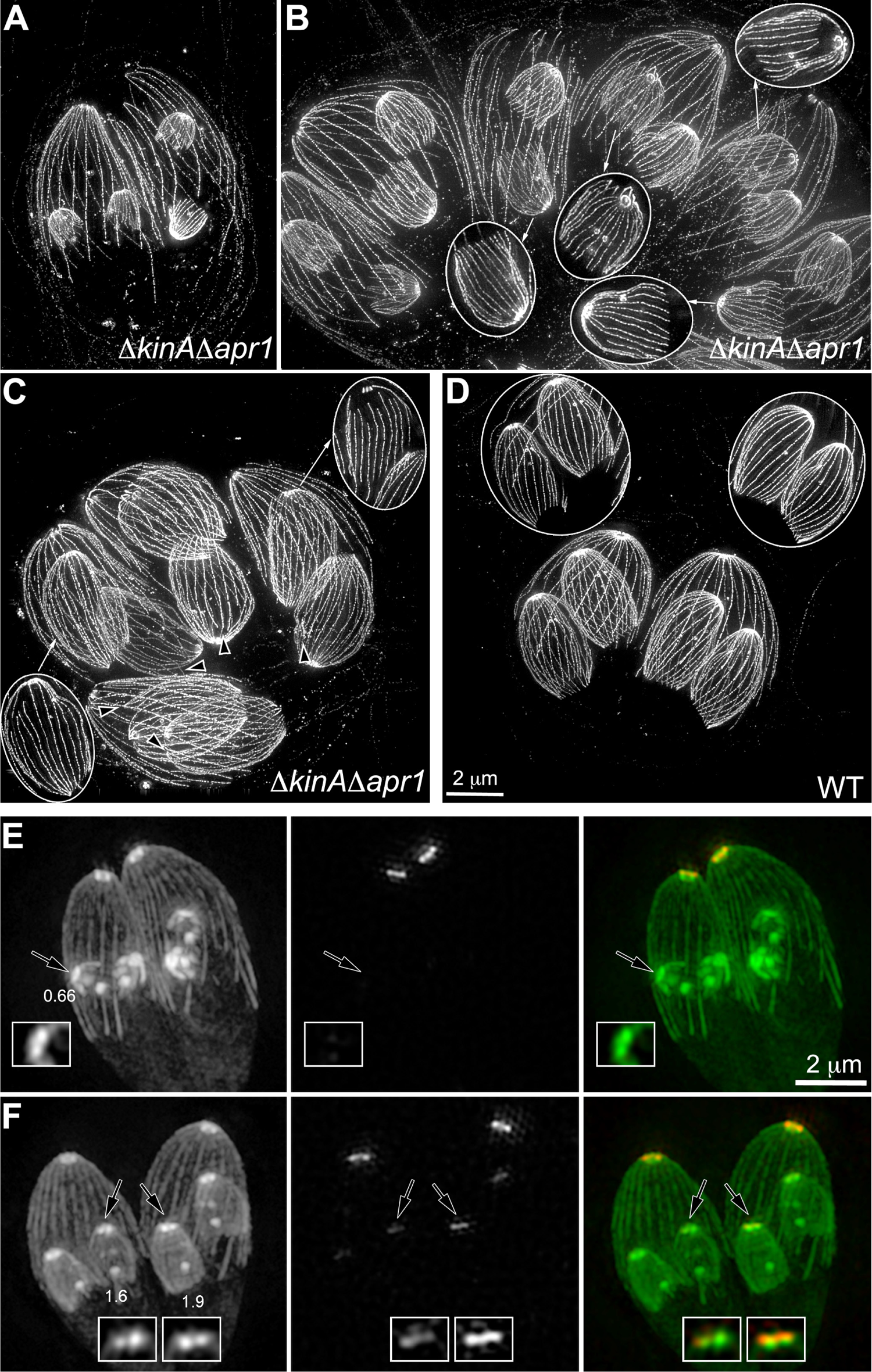
Detachment of cortical MTs is detected in *ΔkinesinAΔapr1* daughters after the MTs are no longer grouped into petals. **A.** Projection of ExM images of *ΔkinesinAΔapr1* parasites with relatively early daughters. Detachment of the cortical MTs is pronounced in mother parasites, but not observed in daughters. **B.** Projection of ExM images of *ΔkinesinAΔapr1* parasites with daughters in which MT detachment can be detected. Insets (1.3X) include the projections of a subset of the sections to show the detachment of the cortical MTs in 4 daughter parasites indicated by arrows. **C.** Projection of ExM images of *ΔkinesinAΔapr1* parasites at a late stage of daughter formation. MTs have detached from the apex in 7 out of 8 daughter parasites. Insets (1X) include the projections of a subset of the sections to show the detachment of the cortical MTs in two daughter parasites indicated by arrows. Black arrowheads indicate clusters of MT detachment in 5 other daughter parasites. **D.** Projection of ExM images of WT parasites at a late stage of daughter formation. Insets (1X) include the projection of a subset of the sections to show the intactness of the daughter MT array. Bars ≈ 2 µm prior to expansion **E-F.** Projections of 3D-SIM images of APR1-mCherry knockin parasites expressing GFP-α1-tubulin. Note that APR1 is not detected in the apical polar ring of early stage daughters where the MT petals are discernible by SIM (E). Insets (2X, contrast enhanced) include the apex of daughters indicated by arrows. The numbers indicate the length of daughter parasites in µm. Image contrast was adjusted in the tubulin images to enable better visualization of structures with weaker signals (*e.g.*, cortical MTs) without oversaturating the brighter signals (*e.g.*, conoid labeling).

To determine when the detachment of the cortical MTs occur in the *ΔkinesinAΔapr1* parasites during later daughter development, we measured the incidence of MT detachment in three groups of daughters that have adopted the dome-like shape but with different lengths of MTs. Using the projections of a subset of daughters viewed perpendicular to their long axis, the length of the MTs for the three groups were estimated to be ∼ 1.2 µm ± 0.02 (mean±s.e.m, n=27), 2.0 µm ± 0.04 (n=28), and 3.6 µm ± 0.1 (n=26), respectively. The percentages of daughters with detached MTs were 4.3±2.6% (mean±s.e.m, n=7 imaging fields, total 70 daughters counted), 18.8±7% (n=9 imaging fields, 52 daughters), and 52.1±4.7% (n=25 imaging fields, 182 daughters), respectively. Therefore MT detachment in the daughters becomes more pronounced as daughter MTs elongate in the *ΔkinesinAΔapr1* parasites (Figure 6A-C). In contrast, MT detachment was observed in only one out of 160 late-stage (3.0 µm ± 0.08, n=52) WT daughter parasites (Figure 6D). The significant lag in MT detachment after array initiation in the *ΔkinesinAΔapr1* mutant suggests that the apical polar ring in normal parasites is reinforced as the MT array develops. Previously we discovered that kinesinA is already present in the daughter apical polar ring at the first sign of daughter formation, but APR1 is not detected until later (Leung *et al*., 2017). Here we examined closely APR1 recruitment to the apical polar ring with respect to daughter growth by measuring APR1-mCherry fluorescence and daughter MT length in live APR1-mCherry knock-in parasites expressing GFP-tubulin. We discovered that the level of APR1-mCherry fluorescence is close to background in daughters shorter than ∼ 1 µm (38±4, mean±s.e.m, arbitrary units, n=111 parasites), 130±6 (n=253) in daughters between ∼1-2 µm, 688±22 (n=167) in daughters between ∼3-4 µm, and 1351 ±14 (n=422) for the mother parasite. Figure 6E-F shows SIM imaging of dividing parasites with daughters of different sizes. Note that in the early stage where the MT petals can still be discerned by SIM (Figure 6E, daughter length ∼ 0.66 µm), APR1-mCherry fluorescence is not visible. By the time that APR1 is clearly recruited to the developing daughter apex, the MT petals are no longer well-defined when imaged with 3D-SIM (Figure 6F), indicating that APR1 recruitment coincides with the symmetry transition of the MT array. We thus propose that the stability of the apical polar ring needs to be strengthened continuously by the recruitment of the later components (*e.g.*, APR1). This is needed to maintain the integrity of the apical polar ring, possibly to counter tension generated at the MT anchoring sites due to daughter expansion, growth, and changing symmetry of the MT array.

## DISCUSSION

The best understood mechanism of the establishment of a polarized MT array comes from studies of animal cells, where the γ-tubulin ring complexes embedded in the centrosome matrix serve to initiate and maintain the polarity of the MT cytoskeleton (Oakley and Oakley, 1989; Stearns *et al*., 1991; Joshi, 1994; Stearns and Kirschner, 1994; Moritz *et al*., 1995; Patel and Stearns, 2002; Kollman *et al*., 2008; Oakley *et al*., 2015). The high concentration of pre-assembled nucleating complexes at the centrosome provides an intrinsic bias in polarity. In addition, by capping the minus end of the microtubules, the effective critical concentration for γ-tubulin nucleated MTs is lower than that for free cytoplasmic MTs, further enhancing the preferential MT growth from the centrosome. During mitosis, the interphase array extensively rearranges and reassembles into the biopolar mitotic spindle, but the core principle of the organization of the MT array remains the same [reviewed in (Mitchison, 1988, 1989; Wadsworth, 1993)]. Another extensively studied MT-based organelle is the flagellum/cilium, which originates from centrioles [reviewed in (Azimzadeh and Marshall, 2010; Bettencourt-Dias, 2013)].

In the asexual stage of *Toxoplasma gondii,* there are five known tubulin-containing structures: the conoid fibers, intra-conoid MTs, cortical MT array, centrioles, and the spindle. During cell division, the spindle is assembled in the nucleus, while the other four tubulin-based structures duplicate (centrioles) or are constructed *de novo* in the cytoplasm (conoid fibers, intra-conoid MTs, cortical MT array). γ-tubulin is found only in the centrosome (Morrissette, 2015; Suvorova *et al*., 2015), which is likely structurally coupled to the mitotic spindle. Therefore, the construction of the other non-centriole tubulin-containing structures has to involve novel components and perhaps novel mechanisms. Our previous work showed that the conoid and cortical MT array are initiated in the vicinity of the centrioles (Hu *et al*., 2006; Hu, 2008; Liu *et al*., 2015), but the resolution was insufficient to discern more structural details.

Here we reveal for the first time the spatial relationship among these structures and how their nascent forms co-develop. Construction of the conoid, intra-conoid MTs, and cortical MTs all begin adjacent to the centrioles with clear structural differentiation in the earliest detectable anlagen (Figure 3-4). It is astonishing that three different supramolecular assemblies of quite different structure are being put together from the same pool of tubulin subunits at the same time within a region of ∼ 500 nm across. The assembly of the conoid and the cortical MT array both proceed towards the centrioles, while the intra-conoid MTs grow within the developing conoid slightly after the nascent conoid first appears as an arc in the end-on view. This indicates that these three sets of structures are initiated through distinct but coordinated mechanisms. This open-sided intermediate of the conoid is intriguing, because it resembles the architecture of the “pseudoconoid” in the free-living, marine relatives of apicomplexans.

The “pseudoconoid” is a half-cone made of a curved sheet of canonical microtubules (Portman *et al*., 2014). The resemblance to the developing conoid is even more suggestive if, as recently proposed, the conoid fibers begin life as canonical microtubules (Li *et al*., 2023) and only later transition to folded ribbons. Many lines of evidence suggest that the ancestor of the apicomplexan parasites descended from a free-living marine protist (Oborník and Lukeš, 2013; Janouškovec *et al*., 2015; Woo *et al*., 2015). One thus has to wonder whether the initiation, development, and maturation of the conoid recapitulates how this structure evolved over hundreds of millions of years.

In the nascent daughters, the cortical MTs are already clustered into petals before the nucleation of all 22 MTs is completed. At a later stage, the 22 MTs are organized into a 5-petaled structure. In WT parasites, both the organization and the number of MTs are essentially invariant from cell to cell. This basic layout of the array is likely determined by the localization and number of the nucleation sites. In adult parasites, the 22 MTs are not clustered, but instead are evenly distributed around the cortex, *i.e.,* arranged with 22-fold rotational symmetry. Therefore the switch from the approximate 5-fold symmetry of the petals to 22-fold symmetry occurs via major rearrangements of MTs rather than addition of new MTs or removal of existing MTs.

Two components of the apical polar ring (kinesinA and APR1) are known to be important for its stability (Leung *et al*., 2017). The loss of these two proteins destabilizes the apical polar ring, but does not affect the nucleation of the MTs in the cortical array. Interestingly, analysis of early daughter structure in this double-knockout mutant shows however that the fidelity of the MT array is affected. Instead of a constant 22 MTs, a significant number of the double knockout mutant makes 23 or 24 MTs. This suggests that 1) the nucleating and scaffolding functions of the apical polar ring components are distinct and 2) the disruption of the scaffold often affects the number of nucleation sites. These observations also highlight the power of the extreme reproducibility of the cytoskeletal structure in *Toxoplasma*, which enables the detection of quite subtle changes in the MT array that would be impossible to detect in most other cell types.

In the adult *ΔkinesinAΔapr1* parasite, the cortical MTs are dispersed from the apex (Leung *et al*., 2017). Here we showed that this detachment occurs in the daughter after the initial MT petals have been established. Incidentally, we found the recruitment of APR1 in WT parasites also accelerates after the initiation stage. This supports the speculation that APR1 reinforces the attachment of the MTs to the developing apical polar ring during the elongation and expansion of the array. Currently, no nucleation factors have been identified for the cortical MT array, conoid, or the intra-conoid MTs. As these structures are distinct from the centrosome, exploring their nucleation mechanisms will reveal new biology. The structural data reported here can serve as guidance for identifying and interpreting the function of nucleating factors and structural components of the organizing centers.

## MATERIALS AND METHODS

### T. gondii strains and host cell cultures

Host cell and *T. gondii* tachyzoite parasite cultures were maintained as described previously (Roos *et al*., 1994; Liu *et al*., 2015; Leung *et al*., 2017; Munera Lopez *et al*., 2022). The *ΔkinesinAΔapr1* parasite was reported in (Leung *et al*., 2017). The *mEmeraldFP-TrxL1* knock-in parasite was reported in (Liu *et al*., 2013). The *APR1-mCherry* knock-in parasite stably expressing eGFP-Tg α1-tubulin (TUBA1) was generated by transfecting a ptubA1-eGFP-TgTUBA1 plasmid into the *APR1-mCherry* knock-in parasite (Leung *et al*., 2017), selected with 20 µM chloramphenicol and cloned. The backbone and CDS arrangement for ptubA1-eGFP-TgTUBA1 are identical with those for ptubA1-mCherryFP-TgTUBA1 (Leung *et al*., 2017).

### Three-dimensional structured-illumination microscopy (3D-SIM)

Cultures of intracellular parasites growing in HFF cells plated in 3.5-cm glass-bottom dishes (MatTek Corporation, CAT# P35G-1.5-21-C) were placed in L15 imaging media [Leibovitz’s L-15 (21083-027, Gibco-Life Technologies, Grand Island, NY) supplemented with 1% (vol/vol) heat-inactivated cosmic calf serum (CCS; SH30087.3; Hyclone, Logan, UT)]. Samples were imaged with a DeltaVision OMX Flex imaging station (GE Healthcare-Applied Precision).

Image stacks for 3D-SIM were collected at z-increments of 0.125 µm using a 60x oil immersion lens (numerical aperture [NA] 1.42) and immersion oil of refractive index 1.520. Images were reconstructed using Softworx (GE Healthcare-Applied Precision) and the point spread function supplied by the manufacturer. Contrast levels were adjusted to optimize the display.

### Sample preparation for Expansion Microscopy (ExM)

The protocol for ExM basically followed the descriptions in (Gambarotto *et al*., 2019; Vincent *et al*., 2022) with slight modifications. *Toxoplasma*-infected monolayers of HFF cells on 12mm round coverslips were fixed with 3.6 % formaldehyde (made from paraformaldehyde) in PBS for 15-30 min at room temperature, washed several times with PBS, and stored at 4 °C in a humidified container. Fixed coverslips were incubated in an anchoring solution (2% acrylamide, 1.4 % formaldehyde, in PBS) at 37 °C in a humidified chamber for 3 - 4 hrs. Prior to gelation, the coverslip with attached cells was incubated with a monomer solution lacking ammonium persulfate (APS) and tetramethylethylenediamine (TEMED) [10% acrylamide, 19% sodium acrylate, 0.1% N,N’-(1,2-dihydroxyethylen)-bisacrylamid (DHEBA)] in the gelation chamber (Vincent *et al*., 2022) on ice for ∼ 15 min. The monomer solution was then drained.

The coverslip was washed briefly with the complete gelation solution (10% acrylamide, 19% sodium acrylate, 0.1% DHEBA, 0.25%TEMED, and 0.25% (w/v) APS in PBS) and then incubated with complete gelation solution in the chamber for 45 min at 37°C. After gelation, the gel with the coverslip was removed from the chamber, placed in a 35 mm dish cell-side up in denaturation buffer (200 mM sodium dodecyl sulfate, 200 mM NaCl, 50mM TrisCl pH 6.8), and shaken until the coverslip detached from the gel. The gel was then placed in a 2 ml Eppendorf tube with 1 ml of denaturation buffer and incubated for 90 min in a heat block at 84 - 85°C. The sample was then washed in dH2O three times before incubation in dH2O at 4°C overnight for full expansion. For immunofluorescence labeling, the fully expanded gel was first washed in PBS briefly 3-4 times and equilibrated in PBS for ∼ 30 min to 1 hr at room temperature. A small piece of the gel was then removed, placed in a glass-bottom dish (MatTek Corporation, CAT# P35G-1.5-21-C) and incubated with primary antibody at 37 °C in a humidified chamber for 3 - 4 hrs, washed in PBS+0.05% tween 20 briefly twice, followed by 15 min washes 3 times. The gel was then incubated with secondary antibodies at 4°C overnight, followed by a 30 to 60 min incubation at 37°C before washing in PBS+0.05% tween 20 as before. The sample was then washed with dH2O three times and left in dH2O for ∼ 1 hr for full re-expansion prior to imaging. For long-term storage, the gel was immersed in PBS at 4°C. Tubulin labeling was carried out with a mouse monoclonal anti-acetylated tubulin antibody (T6793-6-11B-1, Sigma-Aldrich, 1:250) and followed by either a goat anti-mouse IgG Cy3 (115-165-166, Jackson ImmunoResearch Labs, 1:400) or a goat anti-mouse IgG Alexa 488 (A11029, Molecular Probes, 1:400). CEN2 or CEN3 labeling was carried out using rat anti-CEN2 or CEN3 antibodies (1:150 to 1:200) (Leung *et al*., 2017; Munera Lopez *et al*., 2022), followed by either a goat anti-rat IgG Alexa488 (A-11006, Molecular Probes, 1:750) or a goat anti-rat IgG Cy3 (112-165-167, Jackson ImmunoResearch Labs, 1:400). The antibodies were diluted in 2% BSA in PBS.

### Imaging expanded samples

Expanded samples were placed in a MatTek glass-bottom dish. Excess water around the gel was wicked off with filter paper. To maintain humidity in the chamber, wet filter paper strips were plastered against the inner wall of the dish and the dish remained covered throughout imaging. Wide-field image stacks were collected at z-increments of 0.3 µm using a DeltaVision OMX Flex imaging station with a 60x oil immersion lens (NA 1.42) and immersion oil at refractive index 1.520. Images were deconvolved using Softworx (GE Healthcare-Applied Precision) with the point spread function supplied by the manufacturer. Contrast levels were adjusted to optimize the display.

### Determination of the expansion ratio

The degree of expansion was determined by measuring the diameter of the conoid at its basal end in images of the tubulin-antibody labeled ExM samples and in SIM images of live parasites expressing eGFP-tubulin. Image stacks containing parasites viewed perpendicular to their long axis were used for measurement. Within each stack, the optical section closest to the center of the conoid, as judged by the presence in the slice of the intra-conoid microtubules, was selected. The diameter of the conoid at its widest point, i.e., the basal end, was measured using Fiji (Schindelin *et al*., 2012). Measurements were made on 7 SIM images of live parasites (diameter = 0.31 µm±0.01, mean±s.e.m). Measurements on ExM images from three different experiments gave: 1.76 µm±0.02, n= 34; 1.43 µm ±0.01, n= 24; 1.74 µm±0.02, n=41; corresponding to expansion factors of 5.7, 4.6, and 5.6, with a weighted average of 5.4 fold. This is similar to the expansion factors reported in (Gambarotto *et al*., 2019).

Interestingly, the diameter of daughter conoids was consistently ∼10% larger than adult conoids, a small but statistically significant difference (p = 0.0004). It is difficult to measure conoids of daughters in the SIM images, so it is not clear at present whether this difference in size is real or is caused by a difference between adults and daughters in susceptibility to expansion.

### Quantification of APR1-mCherry fluorescence in the parasite apex through daughter development

Live cultures (prepared as noted above for 3D-SIM) were imaged using a DeltaVision OMX Flex imaging station (GE Healthcare-Applied Precision). Wide-field image stacks were collected at z-increments of 0.3 µm using a 60x oil immersion lens (numerical aperture [NA] 1.42) and immersion oil at refractive index 1.520. The same image acquisition parameters (exposure, gain and laser intensity) were used for all images. Quantification of APR1-mCherry fluorescence was carried out using Fiji (Schindelin *et al*., 2012), employing raw images. The corrected fluorescence intensity was calculated by subtracting the background (measured on an area adjacent to the apical polar ring) from the total fluorescence (sum of all pixel values in an oval region enclosing the apical polar ring).

## Acknowledgments

We thank Melissa Molina for tissue culture support, Dr. Jonathan Munera Lopez and Ms Isadonna F. Tengganu for helpful discussions. This study was supported by funding from the National Institutes of Health/National Institute of Allergy and Infectious Diseases (R01-AI132463) awarded to K.H and from the National Science Foundation (2119963) awarded to K.H (co-PI).

